# Melanoma proteomics unravels major differences related to mutational status

**DOI:** 10.1101/198358

**Authors:** Lucía Trilla-Fuertes, Angelo Gámez-Pozo, Guillermo Prado-Vázquez, Andrea Zapater-Moros, Mariana Díaz-Almirón, Claudia Fortes, Rocío López-Vacas, Iván Márquez-Rodas, Ainara Soria, Juan Ángel Fresno Vara, Enrique Espinosa

## Abstract

The aim of the study was to explore the molecular differences between melanoma tumor subtypes, based on BRAF and NRAS mutational status. Fourteen formalin-fixed, paraffin-embedded melanoma samples were analyzed using a high-throughput proteomics approach, coupled with probabilistic graphical models and Flux Balance Analysis, to characterize these differences. Proteomics analyses showed differences in expression of proteins related with fatty acid metabolism, melanogenesis and extracellular space between BRAF mutated and BRAF non-mutated melanoma tumors. Additionally, probabilistic graphical models showed differences between melanoma subgroups at biological processes such as melanogenesis or metabolism. On the other hand, Flux Balance Analysis predicts a higher tumor growth rate in BRAF mutated melanoma samples. In conclusion, differential biological processes between melanomas showing a specific mutational status can be detected using combined proteomics and computational approaches.

## Introduction

Melanoma is the most lethal cutaneous cancer, with over 11,000-15,000 estimated deaths in the United States and Europe every year [1,2]. Better understanding of the molecular biology of this tumor has allowed the development of new effective drugs for the treatment of advanced disease, both in the fields of targeted therapies and immunotherapy [3]. However, as not all patients obtain a benefit from new drugs, further insight into the biology of melanoma is needed.

Gene signatures, genomic hybridization, whole-exome genome sequencing, microRNA analysis and other techniques have widely addressed the genomic landscape of melanoma, contributing to significant advances [4,5]. Given the heterogeneity of melanoma and the complex interaction of this tumor with the immune system, the need for combination of biomarkers assays has been recently proposed to properly analyze the disease [6].

Proteins determine cell phenotype, so proteomics analyses offer the possibility to measure the biologic outcome of cancer-related genomic abnormalities [7]. Mass spectrometry has become the method of choice to assess complex protein samples, and recent technological advances allow the identification of thousands of proteins from tissue amounts compatible with clinical routine. Therefore, proteomics may become a new source of molecular cancer markers offering complementary information to that provided by standard pathology and genomics. We recently demonstrated the feasibility of high-throughput label-free quantitative proteomics to analyze breast cancer from paraffin-embedded samples [8]. In the present study we sought to determine whether high-throughput proteomics combined with computational approaches, such as probabilistic graphical models and Flux Balance Analysis, are useful tools to explore functional differences between groups of melanoma tumors.

## Methods

### Samples

Fourteen melanoma cancer patients were included in the study. FFPE samples were retrieved from Biobanks in IdiPAZ, Hospital Universitario Gregorio Marañón and Hospital Universitario Ramón y Cajal, all integrated in the Spanish Hospital Biobank Network (RetBioH; http://www.redbiobancos.es/). Patients provided informed consent. All experiments were performed in accordance with relevant guidelines and regulations. The histopathological features of each sample were reviewed by an experienced pathologist to confirm diagnosis and tumor content. Eligible samples had to include at least 50% of tumor cells. Approval from the Ethical Committees of Hospital Universitario La Paz was obtained for the conduct of the study.

### Mass-spectrometry analysis protein identification and label-free quantification

Proteins were extracted from FFPE samples as previously described [9]. Peptides were desalted using self-packed C18 stage tips, dried and resolubilized with 15μl of 3% acetonitrile, 0.1% formic acid. Mass spectrometry analysis was performed on a QExactive mass spectrometer coupled to a nano EasyLC 1000 (Thermo Fisher Scientific). Solvent composition at the two channels was 0.1% formic acid for channel A and 0.1% formic acid, 99.9% acetonitrile for channel B. For each sample 3μL of peptides were loaded on a commercial PepMapTM RSLC C18 Snail Column (75 μm × 500 mm, Thermo Fisher Scientific) and eluted at a flow rate of 300 nL/min by a gradient from 2 to 30% B in 85 min, 47% B in 4 min and 98% B in 4 min. Samples were acquired in a randomized order. The mass spectrometer was operated in data-dependent mode (DDA), acquiring a full-scan MS spectra (300-1700 m/z) at a resolution of 70000 at 200 m/z after accumulation to a target value of 3000000, followed by HCD (higher-energy collision dissociation) fragmentation on the twelve most intense signals per cycle. HCD spectra were acquired at a resolution of 35000 using normalized collision energy of 25 and a maximum injection time of 120 ms. The automatic gain control (AGC) was set to 50000 ions. Charge state screening was enabled and singly and unassigned charge states were rejected. Only precursors with intensity above 8300 were selected for MS/MS (2% underfill ratio). Precursor masses previously selected for MS/MS measurement were excluded from further selection for 30 s, and the exclusion window was set at 10 ppm. The samples were acquired using internal lock mass calibration on m/z 371.1010 and 445.1200.

The acquired raw MS data were processed by MaxQuant (version 1.5.2.8), followed by protein identification using the integrated Andromeda search engine. Spectra were searched against a forward Swiss Prot-human database, concatenated to a reversed decoyed fasta database and common protein contaminants (NCBI taxonomy ID9606, release date 2014-05-06). Carbamidomethylation of cysteine was set as fixed modification, while methionine oxidation and N-terminal protein acetylation were set as variable. Enzyme specificity was set to trypsin/P allowing a minimal peptide length of 7 amino acids and a maximum of two missed-cleavages. Precursor and fragment tolerance was set to 10 ppm and 20 ppm, respectively for the initial search. The maximum false discovery rate (FDR) was set to 0.01 for peptides and 0.05 for proteins. Label free quantification was enabled and a 2 minutes window for match between runs was applied. The re-quantify option was selected. For protein abundance the intensity was used, corresponding to the sum of the precursor intensities of all identified peptides for the respective protein group.

Following MS workflow, identified protein groups were filtered by the presence of at least two unique peptides and detectable expression in at least 75% of the samples. Label-free quantification values from these proteins were used for subsequent analyses. Additionally, batch effects were removed using limma package [10] and R v 3.2.5 [11].

### Protein differential expression analyses

Significance Analysis of Microarrays (SAM) was performed using MeV to find significant differences in protein expression among samples [12]. Protein expression patterns were also compared calculating delta values for each biomarker status against the rest of the tumor samples. Proteins showing a change in expression value higher than 1.5 or lower than −1.5 were selected.

### Probabilistic graphical model and activity measurements

R v 3.2.5 [11] and grapHD package [13] were used to build a probabilistic graphical model as previously described [14,15]. The network was split into several branches and Gene Ontology analysis was used to assign a major function to each node. Activity measurements were then calculated by the mean expression of all the proteins related to the assigned node function.

### Flux Balance Analysis

Flux Balance Analysis (FBA) was performed using COBRA Toolbox [16] and whole metabolism human reconstruction Recon 2 [17] both available for MATLAB. As an objective function, biomass reaction supplied by the Recon 2 was used as representative of tumor growth rate. Proteomics expression data was incorporated into the model as described in previous works [15]. Briefly, GPR rules were estimated using the sum for "ORs" expressions and minimum for "ANDs" expressions. Then, E-flux algorithm [18] was used to normalize the GPR values dividing by the maximum value in each tumor and incorporate protein expression data into the model.

### Statistical analyses

GraphPad Prism v6 was used for statistical analyses, whereas Cytoscape was used for network analysis. Gene Ontology Analyses were performed in DAVID webtool selecting “Homo sapiens” background and GOTERM-FAT, Biocarta and KEGG databases.

## Results

### Patients and samples

Primary melanoma samples coming from 14 patients with advanced disease were included. Samples were split into three groups according to mutational status: BRAF-mutant (n=3), NRAS-mutant (n= 5) or double negative (n= 6). BRAF and NRAS mutations had been previously determined in local laboratories with standard polymerase chain reaction-based tests.

### Mass-spectrometry analysis

FFPE melanoma tumor samples were analysed by mass-spectrometry. 4,006 protein groups were identified, of which 1,606 present at least two unique peptides and detectable expression in at least 75% of the samples. Label-free quantification values from these 1,606 proteins were used for subsequent analyses.

### Differential protein expression patterns between subtypes

A Significance Analysis of Microarrays (SAM) was done to find differences among samples at the protein level. Seventeen proteins were found differentially expressed between BRAF mutated and BRAF wild type tumors (Figure 1).

**Figure 1:**
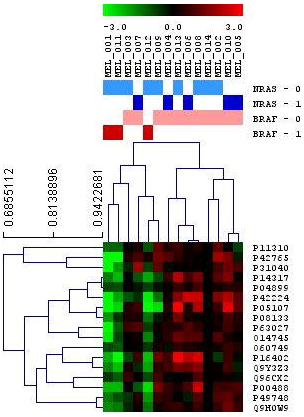
Significance Analysis of Microarrays between BRAF positive and negative tumors.

In addition, delta values between BRAF-mutated and BRAF-wild type, and NRAS-mutated and NRAS-wild type tumors were calculated. Delta values higher than 1.5 or lower than −1.5 were used to perform gene ontology analyses as well. Proteins related with keratinization, epidermis development and cytoskeleton were underexpressed, whereas proteins involved in melanogenesis and extracellular space were overexpressed in BRAF-mutant as compared with BRAF-wild type samples. SAM and delta analyses did not find significant differences between NRAS-mutant and NRAS-wild type tumors.

### Probabilistic graphical model and node activity measurements

A probabilistic graphical model was built using proteomics data without other a priori information. The resulting network was processed to build a functional structure, as described in previous works [14,15]. The resulting network was divided into thirteen branches, and gene ontology analyses were performed to establish functional structure. Finally, twelve principal functions were assigned to different branches and there was a branch to which no main function could be assigned (Figure 2).

**Figure 2:**
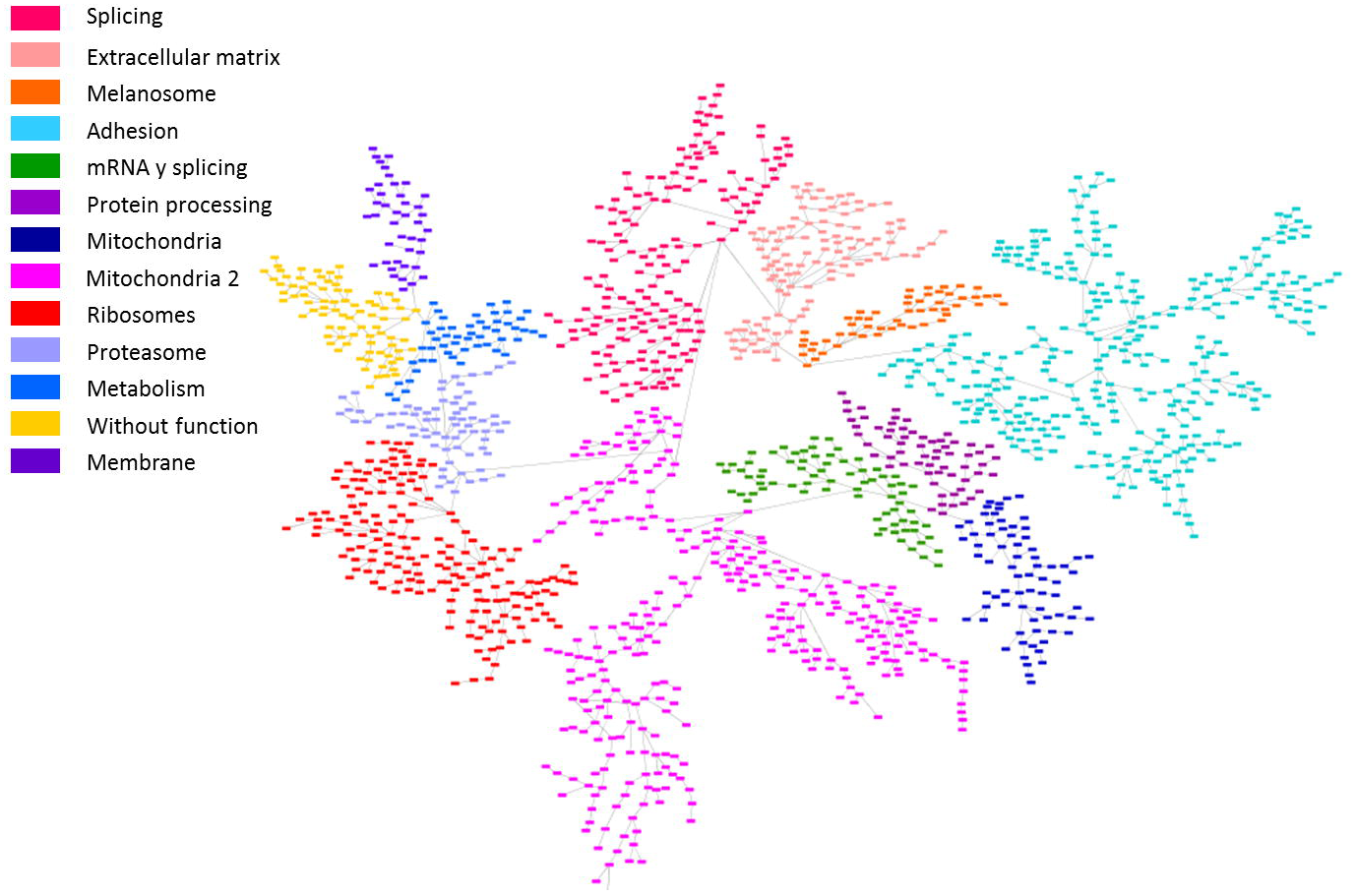
Probabilistic graphical model built using protein expression data from melanoma tumors.

Node activity measurements were calculated for each node using proteins related with the main assigned function and a comparison between BRAF-mutant, NRAS-mutant and double-negative groups was performed. Although the limited number of samples did not allow seeing significant differences, some trends in functional activities were found. For instance, NRAS-mutant had a lower melanosome node activity than BRAF-mutant or double negative tumors. On the other hand, BRAF-mutant tumors had a higher metabolism node activity than NRAS-mutant or double negative (Figure 3).

**Figure 3:**
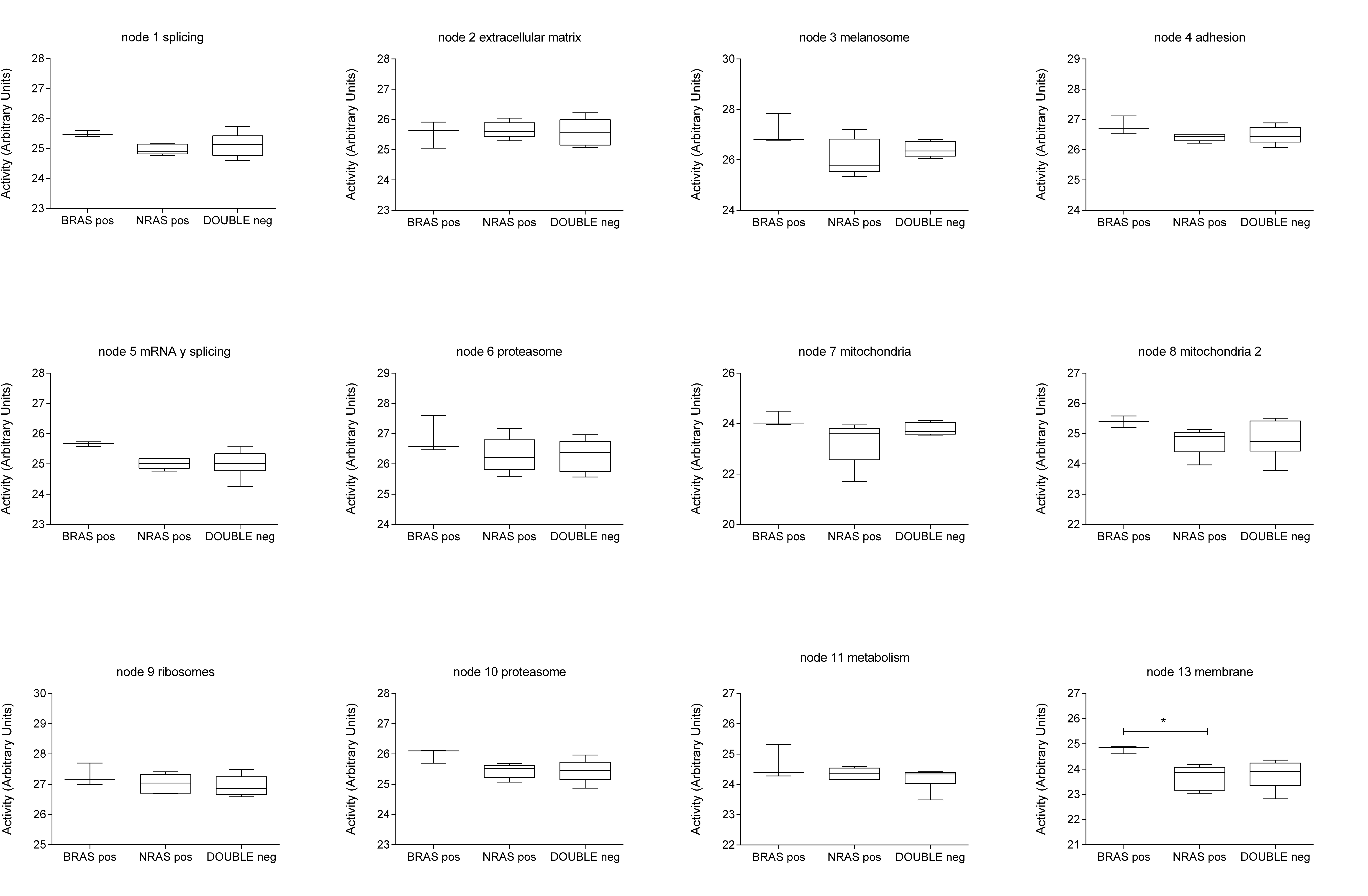
Activity measurements calculated for each network functional node according biomarkers features.

### Flux Balance Analysis

Flux Balance Analysis is a computational approach to assess biochemical networks through the calculation of the flow of metabolites through this network. FBA can be used to calculate the growth rate of an organism or the rate of production of a given metabolite. Our model predicted that BRAF mutated tumors have a higher tumor growth rate than the two other subtypes (Figure 4).

**Figure 4:**
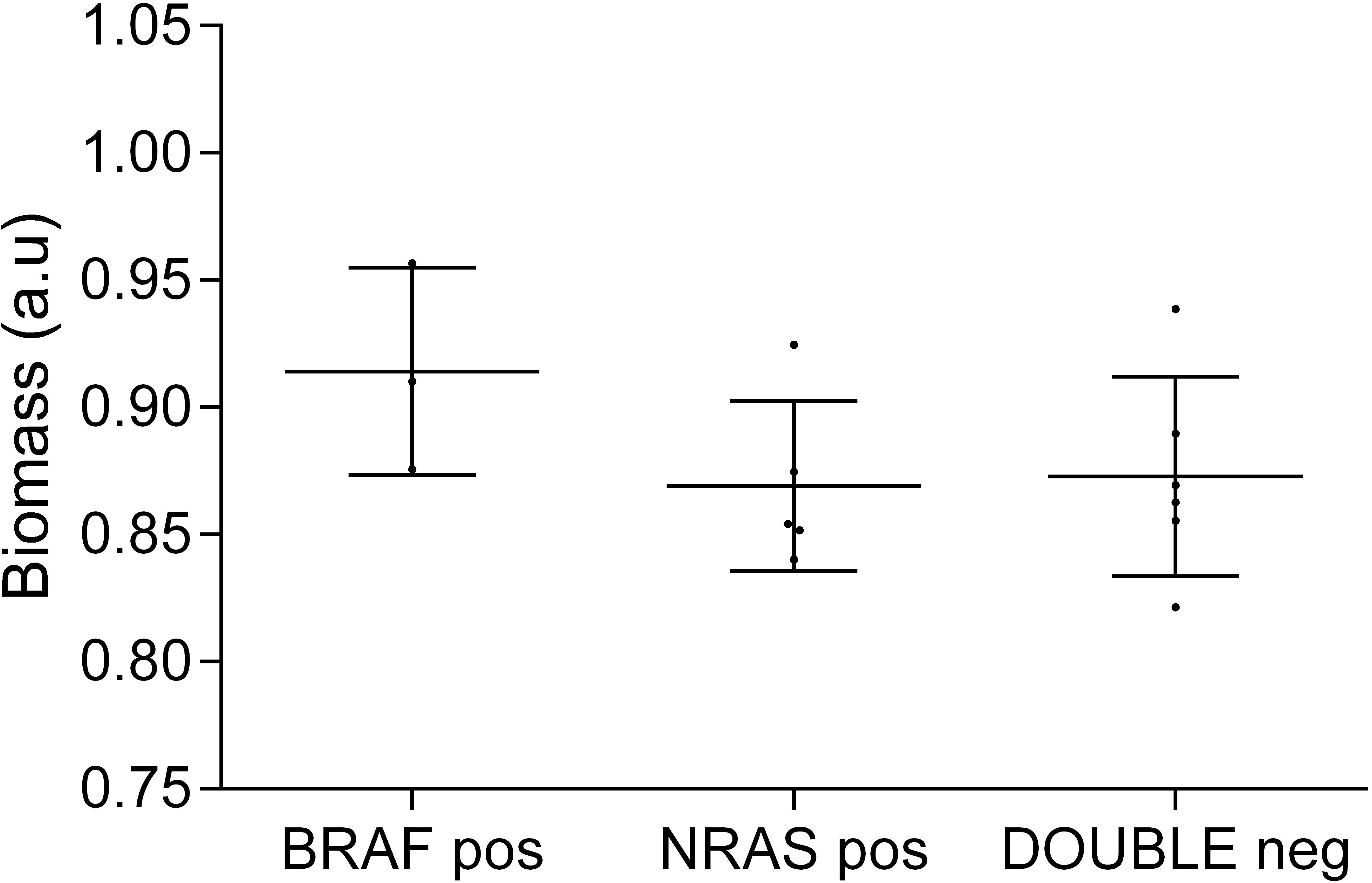
FBA predicted growth rates.

## Discussion

In this study, proteomics coupled with probabilistic graphical models and flux balance analysis were used to characterize differences between melanoma biomarker subgroups in melanoma samples.

Mass-spectrometry workflow allowed the detection of 1,606 proteins with two unique peptides and detectable expression in at least 75% of the samples. Differences in fatty acid metabolism, cytoskeleton or keratinization were observed between BRAF–mutant and BRAF-wild type tumors. Also, differences in functions such as melanogenesis or metabolism were shown between subgroups

SAM and gene ontology analysis found 17 proteins differentially expressed between BRAF-mutant and the two other subgroups (NRAS-mutant and double-negative). These proteins are mainly involved in fatty acid metabolism: acyl-Co A dehydrogenases ACADM, ACAA2 and ACADVL. Another protein with a differential expression is succinate dehydrogenase complex flavoprotein subunit A (SDHA), which encodes a major catalytic subunit of succinate-ubiquinone reductase, a complex of the mitochondria chain, and it was previously related with melanogenesis process [19]. Histidine/aspartate (HD)- domain containing protein 1 (SAMHD1) is implicated in regulation of DNA replication and damage repair and it is proposed to have antiproliferative and tumor suppressive functions in many cancers [20]. Sorting nexin 2 (SNX2) is involved in membrane trafficking of growth factor receptors including epidermal growth factor receptor and c-Met [21]. Another protein differentially expressed is the coagulation factor XIII (F13A1) which it was previously associated with chemotherapy response in melanoma tumors [22]. Potassium channel tetradimerization domain containing 12 (KCTD12) inhibits proliferation in uveal melanoma cells [23]. SLC9A3R1 is involved in suppressing breast cancer cells proliferation [24]. Annexin A6 (ANXA6) acts as a tumor suppressor in skin cancer and it is involved in in the conversion of melanocytes to malignant melanomas [25]. Integrin subunit beta 2 (ITGB2) participates in cell adhesion as well as cell-surface mediated signalling and it is correlated with survival in other cancers such as renal or colorectal tumors [26,27]. It was previously described that metastatic melanoma tumors have a decreased expression of signal transducer and activator of transcription (STAT1) and it could be one of the mechanism by which melanoma can evade immune detection [28]. Finally, G protein subunit alpha i2 (GNAI2) contributes to melanoma cell growth [29].

Differential analyses did not show differences between NRAS-mutant and NRAS-wild type tumors, which are attributable to the small sample size. The present study was limited in this regard because it was designed just as a proof of principle that high-throughput proteomics can be used to study clinical samples of melanoma. Future studies with larger sample size will be needed to establish significant differences among subtypes. Interestingly, it seems that delta analyses and SAM provide complementary information about different protein expression patterns, because differential proteins provided by these two analyses were related to different biological processes.

On the other hand, a probabilistic graphical model was used to generate a network based in protein expression data. It is remarkable that, despite the low number of samples, the probabilistic graphical model clearly showed a functional structure. This type of analysis previously demonstrated its utility to characterize other tumor types such as bladder carcinoma or breast cancer and may complement the information provided by genomics [15]. On the other hand, the high growth rate in BRAF-mutant tumors predicted by FBA agrees with previous knowledge [30].

In summary, our study demonstrates proteomics and computational methods can be applied to the study of clinical melanoma samples. Our results suggest that subgroups defined by mutational status have major differences at protein and functional levels.

## Acknowledgements

We want to particularly acknowledge the patients in this study for their participation and to IdiPAZ, as well as participating Biobanks.

